# Sez6L2 autoimmunity induces cerebellar ataxia in mice

**DOI:** 10.1101/2025.05.28.656724

**Authors:** Carlos J. Reyes-Sepúlveda, Julia Granato, John Randolph, Allison T. Ryan, Alison Hobbins, Angela Stout, Herman Li, Jennetta W. Hammond

## Abstract

Sez6L2 autoantibodies have been reported in patients with subacute cerebellar ataxia, presenting with gait disturbances, frequent falls, slurred speech, and repetitive eye movements. These case studies suggest that autoimmunity against Sez6L2 causes cerebellar damage leading to ataxia. Sez6L2 is a transmembrane protein expressed by most neurons, with the highest levels in the cerebellum. We tested whether immunizing C57BL/6 mice against Sez6L2 could produce an autoimmune response resulting in ataxia symptoms and immune attack on the brain/cerebellum. We found that immunized mice generated significant levels of anti-Sez6L2 antibodies, with IgGs present in the cerebellar parenchyma. Additionally, Sez6L2-immunized mice developed a significant population of Sez6L2-specific T cells targeting two immunodominant epitopes and showed increased CD4^+^ T cell infiltration into the brain. These mice exhibited mild mobility impairments in open field, wire grid walk, and pole test assays. Our results indicate that an autoimmune response to Sez6L2 in mice can lead to mobility impairments and pathology consistent with human cerebellar ataxia associated with Sez6L2 autoantibodies. This new mouse model should be useful for mechanistic studies on this poorly understood autoimmune disease and for pre-clinical testing of therapeutic strategies targeting the immune system in patients with Sez6L2 antibodies.

## INTRODUCTION

Autoimmune diseases of the central nervous system often manifest with symptoms common to movement disorders, such as ataxia (2). Sez6L2 autoantibodies have been reported in patients with subacute cerebellar ataxia accompanied by parkinsonism (3–14). Patients present gait disturbances, frequent falls, slurred speech, and ocular motor symptoms. Some patients also develop cognitive deficits, depression, dizziness, bradykinesia, and/or postural instability. The average age of onset is 59 years (range 35-73) with approximately 70% of affected individuals being female. MRI neuroimaging studies have revealed mild to moderate cerebellar atrophy that worsens over time usually without other inflammatory features visible by MRI. Only about one-third of patients have reported abnormal CSF results suggestive of inflammation. However, Sez6L2 autoantibodies have been found in both the CSF and serum. Screening for malignant tumors suggests a paraneoplastic etiology in about 25% of patients. Response to immunotherapy has been variable, with at least half of patients exhibiting mild improvement and/or stabilization of symptoms with treatment (3–14). However, some patients may have received treatment too late or for too short a period to accurately assess efficacy.

Sez6L2 belongs to the Sez6 family, which includes Sez6, Sez6L, and Sez6L2. They are transmembrane proteins predominantly expressed by neurons. Although the family shares a similar domain structure, they are only about 50% identical in amino acid sequence. All Sez6 family members can be cleaved near the transmembrane region by BACE enzymes, releasing their extracellular domains into the CSF (15, 16). In experimental models, the Sez6 family has been shown to modulate synapse numbers, synaptic plasticity, and dendritic morphology in the cortex and hippocampus, as well as neuronal connectivity in the cerebellum (17–20). Genetic knockout of all Sez6 family genes in mice results in impaired motor functions, motor learning, and cognition (17, 18). The phenotype of mice with a genetic loss of only Sez6L2 has not yet been reported in the literature.

Sez6L2 is genetically linked to autoimmune conditions such as multiple sclerosis and systemic lupus erythematosus, as well as Alzheimer’s disease, autism spectrum disorder, and other neurodevelopmental conditions (21–32). Additionally, Sez6L2 peptides in cerebrospinal fluid have been identified in several biomarker screens for multiple sclerosis, epilepsy, and Alzheimer’s disease (33–35). While the molecular function of Sez6L2 remains unclear, our lab has discovered one mechanism for Sez6L2 as an inhibitor of the complement cascade (36). It has also been proposed to serve as a trafficking/sorting receptor or binding partner of GluR1 receptors and adducin(5, 37–39).

To test the hypothesis that Sez6L2 autoimmunity damages the cerebellum leading to cerebellar ataxia, we immunized C57BL6 mice with Sez6L2 and investigated the autoimmune response. Our findings indicate that autoimmunity against Sez6L2 in mice results in a mild motor impairment phenotype, resembling the human condition of late-onset cerebellar ataxia with associated-Sez6L2 autoantibodies. Mice immunized with Sez6L2 generated significant levels of anti-Sez6L2 antibodies and a significant population of Sez6L2-specific T cells. Additionally, these mice exhibited increased infiltration of CD4^+^ T cells and IgG into the brain or cerebellar parenchyma. This new mouse model should prove valuable for mechanistic studies of this poorly understood autoimmune disease and for pre-clinical testing of therapeutic strategies targeting the immune system in patients with Sez6L2 antibodies.

## MATERIALS AND METHODS

### Sez6L2 Recombinant Protein Preparations

The extracellular domain (1–742) of human Sez6L2 (GenBank: BC000567; Protein ID: AAH00567.1; encoding the splice variant that is 809 amino acids in length) was cloned into the pEZYmyc-HIS vector with a C-terminal Myc and 6xHis tag. Likewise, the extracellular domain (1–846) of mouse Sez6L2 (NM_144926.5; Protein ID: NP_659175.1; encoding the splice variant that is 923 amino acids in length) was cloned into a mammalian cloning vector with a CAG promoter and C-terminal 10xHis tag.

HEK293 cells were transiently transfected with the Sez6L2 expression vectors and recombinant Sez6L2 proteins were purified from the media in two separate rounds after 48 and 96 hours. Media was collected, filtered through a 0.2uM filter, supplemented with protease inhibitor cocktail (Sigma Aldrich, P8849; 1:1000), and run through a column with Ni-NTA agarose beads (Thermo Fisher Scientific, R90115). Columns were washed with PBS containing 25 mM imidazole and eluted in PBS with 250 mM imidazole. The elution was then desalted and concentrated through multiple rounds of centrifugation and resuspension in PBS using an Amicon Ultra centrifugal filter with 30 kDa MWCO (MilliporeSigma; UFC903024). Protein concentration was determined by absorbance at 280nm using a NanoDrop Spectrophotometer (ThermoScientific).

### Basic Animal Care

Male and female C57BL6/J mice were obtained from the Jackson Laboratory (Stock #000664) and were immunized between 12 to 15 weeks of age. Mice were maintained on a 12-hour light/dark cycle, with ad libitum access to food and water. Mice were group housed with 3-5 mice per cage. Mice were randomly assigned to immunization groups and housed in mixed group cages. Experimenters were blind to the experimental groups until the final stages of data analysis. All animal procedures and animal behavior assessments were performed during the light cycle. Before each procedure or behavior session, animals were habituated to the testing room for 1 hour. For behavior analysis, males were tested before females.

### Sez6L2 Immunization

Recombinant human or mouse Sez6L2 protein was mixed in PBS with Complete Freund’s Adjuvant (1:1 dilution, Sigma-Aldrich, F5881) for a final concentration of 5 mg of Sez6L2 per mL of emulsion. The Freund’s Adjuvant was supplemented with non-viable, desiccated *Mycobacterium tuberculosis* (Becton, Dickinson, and Company, 231141) with a final concentration of 2 mg/mL of emulsion. Mice were immunized subcutaneously with a total of 200ul of emulsion (containing 1 mg Sez6L2, and 400ug *Mycobacterium tuberculosis)* at two or four sites over the upper and lower back. Sham-immunized controls were injected with a similar control emulsion containing no recombinant Sez6L2 protein. All mice received intraperitoneal injections of pertussis toxin (250ng, Hooke Laboratory, BT-0105) in saline solution, on 0- and 1-day post-immunization (dpi). Mice received a booster immunization of 200µg human or mouse recombinant Sez6L2 in incomplete Freund’s adjuvant (Sigma-Aldrich, 344291; 100ul of emulsion with 1 mg/mL Sez6L2), followed by administration of 250ng of pertussis toxin again on day 0 and 1 post-boost. Mice that developed injection site reactions (mild ulcerative dermatitis) were treated topically with Vetericyn (hypochlorous solution (HOCl). For cohort 1, we immunized 47 mice and tested 46 total mice. One sham mouse was removed from the study as it developed a rectal prolapse. Cohort 1 consisted of: Sham N=10 male, 11 Female (21 total); H-Sez6L2: N=9 male, 8 female (17 total); M-Sez6L2 N=4 male, 4 female (8 total). For cohort 2, we immunized 19 mice: Sham N=6 male, 3 female (9 total); H-Sez6L2 N=6 male, 4 female (10 total).

### Behavioral Assays

#### Open Field

Mice were placed in an empty, open-top chamber box for 10 minutes while being recorded by an overhead camera and analyzed using the video tracking software, ANY-maze (Stoelting Co.).

### Wire Grid/Foot Fall Test

Mice were placed on a plastic-coated wire grid for 2-5 minutes while being recorded by a camera in a horizontal position (viewing both above and below the wire grid). Videos were analyzed manually for the number of times a foot fell between the wires with at least half the foot beneath the wire surface (i.e. foot faults) and the amount of time (in minutes) the mouse was moving. Grooming and exploring while standing were not counted as time mobile. Data was calculated as foot faults per minute of mobile activity. If a mouse failed to move on the grid at least 10 seconds per minute, they were excluded from the foot fault study. Two mice from cohort 2 (1 sham and 1 H-Sez6L2 immunized) were excluded due to limited movement on the wire grid.

### Pole Test

Mice are placed facing downward at the top of a 55-60 cm vertical metal pole with a diameter of ∼1 cm and small grooves to facilitate easy gripping. The test measures strength and coordination by assessing the time it takes the animal to descend to the cage floor. Mice receive 3 training runs followed by 3-4 recorded trials. If the animal fell, the maximum duration of 20 seconds was assigned for that trial.

### Rotarod

Mice received three training sessions on the rotarod (Panlab, Harvard Apparatus, 76-0770) lasting three minutes each at a fixed rotation speed of 5 rotations per minute (RPM). During this training, mice were put back on the rod immediately if they fell off so they all experienced 3 minutes of training on the rotarod. The following day mice received one final training session lasting 1 minute at 5 RPM. Mice were then tested in 4 separate trials using the accelerating mode set to increase from 4 to 40 RPM over 5 minutes with an inter-trial interval of at least 10 minutes. The time each mouse stayed on the rotarod was recorded as the latency to fall and the best three trials were averaged for each mouse.

### *Isolation of* Splenocytes and Lymph Node Cells

Mice were anesthetized with ketamine and xylazine followed by cardiac perfusion with PBS. For some experiments, the spleen and two inguinal lymph nodes were dissected and processed together. In other experiments, only the spleen was collected and processed. The spleen and lymph nodes in RPMI media were mashed and passed through a 70 µm nylon filter (pluriSelect, 43-50070-51). Cells were then pelleted (400xg for 10 mins) and resuspended in ACK Lysis Buffer (Gibco, A1049201). After a 2–5- minute incubation, cells were pelleted again and resuspended in serum-free media (Cellular Technology Limited, CTLT-010) with 2 mM L-glutamine and 100U/mL Penicillin-Streptomycin (PenStrep; Gibco, 15140122, used 1:100). This supplemented serum-free media is referred to simply as serum-free media in the ELISPOT and *in vitro* stimulation with Sez6L2 sections below.

### IFNγ ELISPOT Assay with recombinant Sez6L2 protein

Mixed splenocyte and lymph node cells were plated on IFNγ ELISPOT pre-coated plates (Cellular Technology Limited, mIFNgp-2M) in serum-free media at a cell density of 200,000 cells per well with or without 8 μg/mL mouse or human recombinant Sez6L2 protein. The cells were incubated at 37°C for 24 hours and processed according to the ELISPOT kit instructions. Briefly, plates were washed twice with PBS and twice with PBSt (PBS with 0.05% Tween). Then, plates were incubated for two hours with a biotinylated anti-mouse IFNɣ detection solution (1:1000). Plates were then washed three times with PBSt and incubated for 30 minutes at room temperature with Strep-AP (1:1000). Plates were washed twice with PBSt and twice with distilled H_2_O. Plates were then incubated for 15 minutes at room temperature with Blue Developer Solution followed by three rinses in H_2_O. Plates were then air dried for 24 hours and IFNɣ+ spots were counted using an ELISPOT reader (ImmunoSpot 5.2.12; Cellular Technology Limited).

### Sez6L2 Peptide Library ELISPOT

A 15mer peptide library with a 10 amino acid overlap that contained peptides covering the extracellular regions of both the human and mouse Sez6L2 proteins was synthesized as a Cleaved PepSet by MIMOTOPES with the yield of each peptide estimated at 1mg. Peptides were dissolved in DMSO at a concentration of 2mg/mL and diluted to a final working concentration in serum-free media of 10ug peptide/mL and 0.5% DMSO. Splenocytes from individual mice (5 immunized, 3 sham) were plated at a density of 350,000 cells per well on BD Mouse IFNγ ELISPOT Set kit (BD Biosciences, 551951) plates coated with anti-IFNγ antibody (1:1000). Cells were incubated with each of the 188 peptides in the library (1 per well) or full-length recombinant mouse Sez6L2 protein for 24 hours at 37°C. Plates were processed according to kit instructions, which were similar to the protocol described above. Spots were developed with HRP/AEC Substrate (BD Biosciences, 551951) and counted using an ELISPOT Reader (ImmunoSpot S6 Macro analyzer; Cellular Technology Limited). Finally, we compared the immunodominant peptides identified by this assay to predictions made by the Immune Epitope Database (IEDB). MHCII binding predictions for the C57BL6 MHCII allele H2-IAb were made on 3/15/2024 using the IEDB analysis resource NetMHCIIpan (ver. 4.1) tool (40).

### *In Vitro* Stimulation and Flow Cytometry Analysis of Splenocytes and Lymph Node Cells

Mixed splenocyte and lymph node cells were plated at two million cells per well in a U-bottom 96 well plate with 200 μL serum-free media supplemented with 2 mM L-glutamine and with or without 10 μg/mL mouse recombinant Sez6L2 protein. The cells were incubated at 37°C for either 18 or 68 hours. Then cells were treated with Brefeldin A (BioLegend, 420601; 1:1000) and incubated for another 6 hours. All following steps were done on ice, with pre-chilled buffers, and protected from light. Cells were pelleted (spin 400xg for 5 minutes) and stained with Ghost Dye Violet 450 (Tonbo Biosciences, 13- 0863-T100) in PBS for 10 to 15 minutes. Plates were washed once in FACS buffer (PBS with 0.5% BSA) and then incubated for 10 minutes with an Fc block (1:25 dilution, BioLegend, Fc Block - Mouse TruStain FcX™ PLUS, 156604). Cell surface antigens were then stained for 30 minutes using antibodies to CD3 (Super Bright Violet 570, Bio-Rad, MCA500SBV570), CD4 (AlexaFluor488, BioLegend, 100532), CD8a (PE- Vio770, Miltenyi Biotechnology, 130-120-817), CD14 (Super Bright 436, Thermo Fisher, 62-0141-80) and/or CD19 (Violet Fluor 450, Tonbo Biosciences, 75-0193-U025). Cells were then washed three times with FACS buffer and incubated in FluoroFix^TM^ buffer (BioLegend, 422101) for 60 minutes. The cells were resuspended in FACS buffer and incubated at 4°C overnight. The following day, the cells were permeabilized by incubating for 5 minutes in Intracellular Perm Buffer (BioLegend, 421002). The cells were washed two more times in Intracellular Perm Buffer then stained for 30 minutes with antibodies for IFNɣ (APC, Tonbo Biosciences, 20-7311-U025), TNF-α (Brilliant Violet 785, BioLegend, 115543), IL-2 (Alexa Fluor 700, BioLegend, 503818), IL-17a (Brilliant Violet 650, BioLegend, 506930), and/or IL-4 (PE-Dazzle 594, Biolegend, 504131). Next, cells were washed three times in FACs buffer and analyzed using a BD LSR II flow cytometer. Data was analyzed by FlowJo^TM^ Software (Version 10.8.2: BD Life Sciences).

### Isolation and Flow Cytometry analysis of CD45^+^ cells from the brain

Mice were anesthetized with ketamine and xylazine (100 and 10 mg/kg, respectively), followed by cardiac perfusion with PBS. Dissected brains were minced and then incubated for 30 minutes at 37°C in 3 mL RPMI with 0.75-1.1 mg/ml Collagenase D (Roche, 11088866001), 100 μg/mL DNAse (Roche 11088866001), 5% FBS, and 10 mM HEPES). Tissue was minced and collected into a centrifuge tube and incubated for 30 minutes at 37°C with 125 µL of collagenase D (Roche Diagnostics, 11088866001) and 30 µL of DNase 1 (100 μg/mL, Roche Diagnostics, 11284932001). Then EDTA was added to final concentration of 2.5 mM and all following steps were performed on ice. The brain slurry was passed through a 70 µm nylon filter (pluriSelect, 43-50070-51), pelleted by spinning at 400xg for 7 minutes, and then resuspended in 38% Percoll (Sigma Aldrich, P1644). The Percoll solution was centrifuged for 30 minutes at 500xg at room temperature with no break. Then myelin was removed from the top along with most of the supernatant. The cell pellet was resuspended in the remaining 100-200µL of supernatant and transferred to a clean 15 mL conical tube with FACS buffer containing 1mM EDTA and 25 μg/mL DNase. Cells were pelleted by spinning at 400xg for 10 minutes, then resuspended in FACS buffer and transferred to a v-bottom 96 well plate for cell surface antigen staining following the same basic protocol outlined above for the splenocyte/lymph node cells but using the following antibodies/markers: CD45 (APC-ef780, Fisher Scientific, 50-112-9642), CD4 (AlexaFluor488, BioLegend, 100529), CD8 (PE, BioLegend, 100707), CD19 (BV785, BioLegend, 115543), CD11b (PE-efluoro610, Fisher Scientific, 50-112-9079), Ly6G (BV605, BD Bioscience, 563005), Ly6C (AlexaFluor647, BioLegend 128009), CD11c (PE-Cy5, BioLegend, 117316), and Ghost Dye Violet 510 (Tonbo Bioscience, 13-0870-T100). After a 30-minute incubation with antibodies, cells were washed two times with FACS buffer and then fixed by incubating them in 2% PFA in PBS for 20 minutes at 4°C. Cells were washed and resuspended with FACS buffer, stored overnight at 4°C, then analyzed using a BD LSR II flow cytometer. Data was analyzed by FlowJo^TM^ Software.

### Sez6L2 Antibody ELISAs

An enzyme-linked immunosorbent assay (ELISA) was used to measure Sez6L2 antibody concentrations in mouse serum, collected at three- and six-week post-immunization. ELISA plates (Nunc, MaxiSorp, 96 well plates, 44-2404-21) were coated with recombinant mouse Sez6L2 at 1ng/µL in 100µL PBS for ∼12-18 hours at room temperature. Wells were then washed three times with PBSt and blocked with 1% BSA in PBS for 30-60 minutes. Mouse serum was diluted in PBS (1:1000) then added to the plate in duplicates and incubated at room temperature for 60 minutes. Wells were again washed three to four times with PBSt and incubated with a goat HRP-conjugated anti-mouse IgG (Bio-Rad, 1706516, 1:15,000) for 60 minutes at room temperature. Wells were then washed three to four times with PBSt prior to colorimetric development using TMB substrate (ThermoScientific, N301) and a 0.18 M sulfuric acid stop solution. Absorbance at 450 nm was measured using a SpectraMax M5 plate reader. Antibody levels were normalized across ELISA plates using serum from one H-Sez6L2 immunized mouse that was measured in all assays.

### Immunohistochemistry (IHC) and Imaging

Mice were anesthetized with ketamine/xylazine (100 and 10 mg/kg, respectively) and cardiac perfused with PBS containing EDTA (1.5 mg/ml) followed by 4% paraformaldehyde (PFA) in PBS. Brains were post-fixed for 24 hours in 4% PFA, then rinsed and stored in PBS at 4°C. The cerebellum and underlying brainstem were isolated from the rest of the brain, incubated in 30% sucrose until the tissue fell to the bottom of the tube, followed by embedding and freezing in OCT compound. The cerebellum was cut into 40 μm-thick sagittal sections using a cryostat and sections were stored in a cryoprotectant mixture of 30% PEG300, 30% glycerol, 20% 0.1 M phosphate buffer, and 20% ddH_2_O at −20°C.

IHC was performed on free-floating sections. The sections were rinsed and incubated in PBS until sections were free from the OCT and cryoprotectant. Then sections were incubated in 100 mM glycine in PBS for 30 minutes followed by two PBS washes and incubation for 30 minutes in blocking buffer (consisting of 1.5% BSA, 3% normal donkey serum (Jackson Immunoresearch Laboratories, 017- 000-121), 0.5% Triton X-100 and 1.8% NaCl in PBS). The primary antibody, IBA1 (Wako, 019-19741, rabbit polyclonal; used at 1:500), was diluted in blocking buffer and incubated with brain sections for 2 days at room temperature with agitation. Sections were then washed three times for 30 minutes in 1× PBS with 1.8% NaCl and then incubated overnight at room temperature with secondary antibodies in blocking buffer (Alexa 488 Donkey anti-mouse F(ab’)₂; Jackson ImmunoResearch Laboratories, 715-546- 151; used at 1:400 to stain endogenous IgGs in the cerebellum; and Rhodamine Red™-X Donkey Anti-Rabbit F(ab’)₂, 711-296-153, used at 1:100). Finally, sections were washed twice with PBS with 1.8% NaCl and stained with DAPI (ThermoScientific, D1306). Sections were washed twice more in PBS and mounted on slides with Prolong Diamond antifade mountant (ThermoFisher Scientific; P36961) and coverslips. IHC sections were imaged with a Leica Stellaris 5, Laser scanning confocal microscope with a white light laser. FIJI software was used for image analysis and image preparation for figures. Image stacks are displayed as maximum intensity projections.

### Statistics

GraphPad Prism software version 10.3.1 for Windows (La Jolla California USA) was used to perform all statistics. Statistical tests and N values for each experiment are specified in the figure legends. Experimental N’s always refer to individual mice. We defined significance as *p <*0.05 and used the following markings on graphs * p<0.05, ** p<0.01 ***, p<0.001, and ****p<0.0001. All data are expressed as the mean ± standard error of the mean (SEM).

## RESULTS

Sez6L2 is a transmembrane protein expressed by most neurons throughout the brain (17, 18). It’s highest expression levels occur in the cerebellum (Figure 1D; GTex Portal; (17, 18)), where it is prominently expressed by Purkinje cells and localizes throughout the soma and dendritic arbors (Figure 1A). It is also expressed by granule cells, Golgi cells, interneurons, and oligodendrocytes (Figure 1 A, C). Immunostaining of non-permeabilized cultured hippocampal neurons shows that a significant portion of Sez6L2 is localized to the neuronal cell surface (Figure 1B).

**Figure 1:**
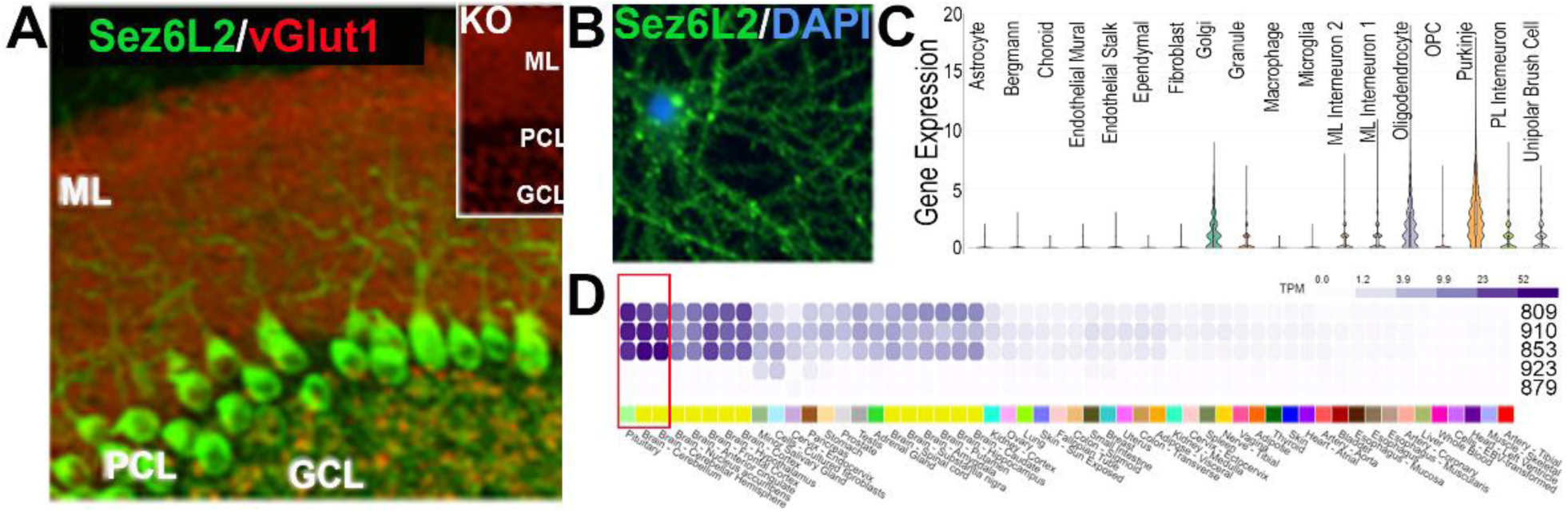
Sez6L2 is expressed by neurons throughout the brain with the highest expression in the cerebellum. A) IHC of Sez6L2 and vGlut1 in the mouse cerebellum. Insert shows no Sez6L2 staining in Sez6L2 knockout mice. ML = molecular layer, PCL = Purkinje cell layer; GCL = granule cell layer. B) Sez6L2 is localized to the cell surface of cultured hippocampal neurons (Sez6L2 antibodies stain non-permeabilized cells.) C) scRNAseq data from adult mouse cerebellum shows Sez6L2 expression in all neurons of the cerebellum and oligodendrocytes (data from Broad Institute Singe Cell Portal viewer of dataset published in (1)). D) Human Sez6L2 expression across all tissues shows Sez6L2 is primarily limited to the nervous system (color coded yellow) with the highest expression in the cerebellum and pituitary (highlighted by the red box). The main transcripts expressed encode Sez6L2 proteins that are 809, 910, and 853 amino acids in length. Image was modified from GTex Portal Transcript Browser.

To generate a model of Sez6L2 immunity, we immunized male and female C57BL/6J mice with the full extracellular domain of human (H-)Sez6L2 or mouse (M-)Sez6L2 in complete Freund’s adjuvant and boosted them three weeks later with Sez6L2 in incomplete Freund’s Adjuvant. Sham immunized mice received similar adjuvant without any protein. Pertussis toxin was given to all groups at days 0 and 1 and following the booster dose. Human Sez6L2 is 95% identical to mouse Sez6L2. We tested both the human and mouse proteins because we hypothesized that, like other autoimmune models (41), the few unique amino acids in the human Sez6L2 protein antigen may help bypass tolerance mechanisms to generate an autoimmune reaction targeting the endogenous mouse Sez6L2.

We found that mice immunized with the human Sez6L2 protein developed a moderate motor- impairment (Figure 2). Specifically, when tested in an open field assay at 2.5- and 5.5-weeks post immunization, the H-Sez6L2 immunized mice traveled less distance and spent more time immobile than sham immunized mice (Figure 2 A, B). The M-Sez6L2 immunized mice were not significantly different from sham immunized mice. On a wire grid/foot fault test that assesses motor function and limb coordination, the H-Sez6L2 immunized mice had significantly more foot faults compared to sham immunized mice (Figure 2 B). Although the M-Sez6L2 immunized mice did not reach statistical significance, they also showed a trend for increased foot faults. Male H-Sez6L2 (and to a lesser extent male M-Sez6L2) immunized mice exhibited some minor weight loss compared to sham controls (Figure 2 C). The weight of female mice was unaffected by the immunizations. Despite the sex-differences in weight loss, both male and female mice displayed motor impairments upon immunization with Sez6L2(see Supplemental Figure 1) and two-way ANOVAs found no significant differences in the motor deficits due to sex. These findings establish that immunization with human Sez6L2 induces a moderate ataxia phenotype in mice, providing a robust model for studying the immune mechanisms underlying Sez6L2 autoimmunity and its associated neurological deficits.

**Figure 2:**
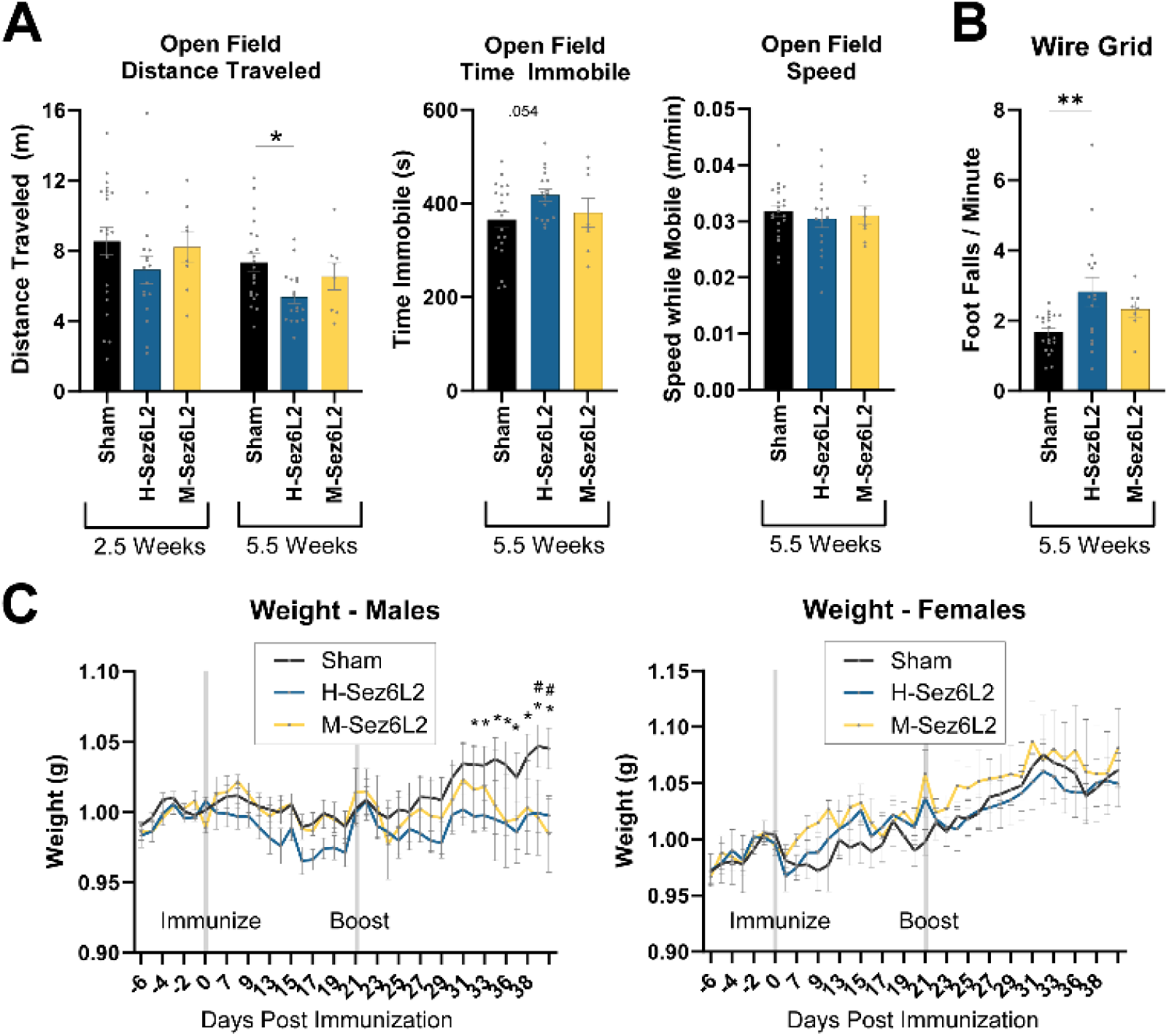
Mice immunized with recombinant human Sez6L2 protein develop significant motor impairments. A) 10-minute open field assay performed at 2.5- and 5.5-weeks post-immunization on a mixed-sex cohort. Graphs show distance traveled, time immobile, and average speed while moving. Statistics: Distance and Speed: 1-way ANOVAs with Tukey’s MCTs, Time Immobile: Welch’s ANOVA with Dunnet’s T3 MCT. B) Foot faults per minute of mobile activity on a wire grid performed at 5.5 weeks post-immunization. 1-way ANOVA with Tukey’s MCT. C) Male and female weight changes post immunization (normalized to day 0). Grey horizontal bars indicate the initial immunization at day 0 and the booster immunization at day 21. 2-way ANOVA with Dunnet’s MCT. * p<0.05 for sham vs H-Sez6L2; # p<0.05 for sham vs M-Sez6L2. Sham N=10 male, 11 Female; H-Sez6L2: N=9 male, 8 female; M-Sez6L2 N=4 male, 4 female.

Next, we assessed the generation of autoreactive T cells against mouse Sez6L2. Using an IFNγ ELISPOT assay, we found that H-Sez6L2 and M-Sez6L2-immunized mice both generated significant populations of Sez6L2-specific T cells (Figure 3 A, B). Building upon these data, we next analyzed autoreactive CD4^+^ and CD8^+^ T cells directly using flow cytometry. After stimulation of combined splenocyte/lymph node cells from Sez6L2- or sham-immunized mice with recombinant mouse Sez6L2 protein for 24 or 72 hours, there were significant or near significant populations of CD4^+^ T expressing IFNɣ, TNFα, IL-4, or IL-17 from our both H- and M-Sez6L2 immunized mice compared to shams (Figure 3 C, D). This suggests the Sez6L2 immunization model generated multiple CD4^+^ T cell subtypes (specifically: Th1, Th2, and Th17) capable of targeting endogenous Sez6L2. Additionally, significant populations of IFNɣ+ CD8^+^ T cells were also found in the Sez6L2-stimulated splenocyte/lymph node culture. This may indicate that a robust pool of Sez6L2-specific CD8^+^ T cells capable of targeting endogenous Sez6L2 were generated in Sez6L2 immunized mice compared to sham immunized mice (Figure 3 E). However, because these “Sez6L2-specific” CD8^+^ T cells were tested in mixed splenocyte cultures with recombinant protein stimulation, they could be the result of cross-activation by the experimental milieu and a more targeted experimental approach is still necessary in the future to fully assess the generation of Sez6L2-specific CD8^+^ T cells and their potential cytotoxic functions.

**Figure 3:**
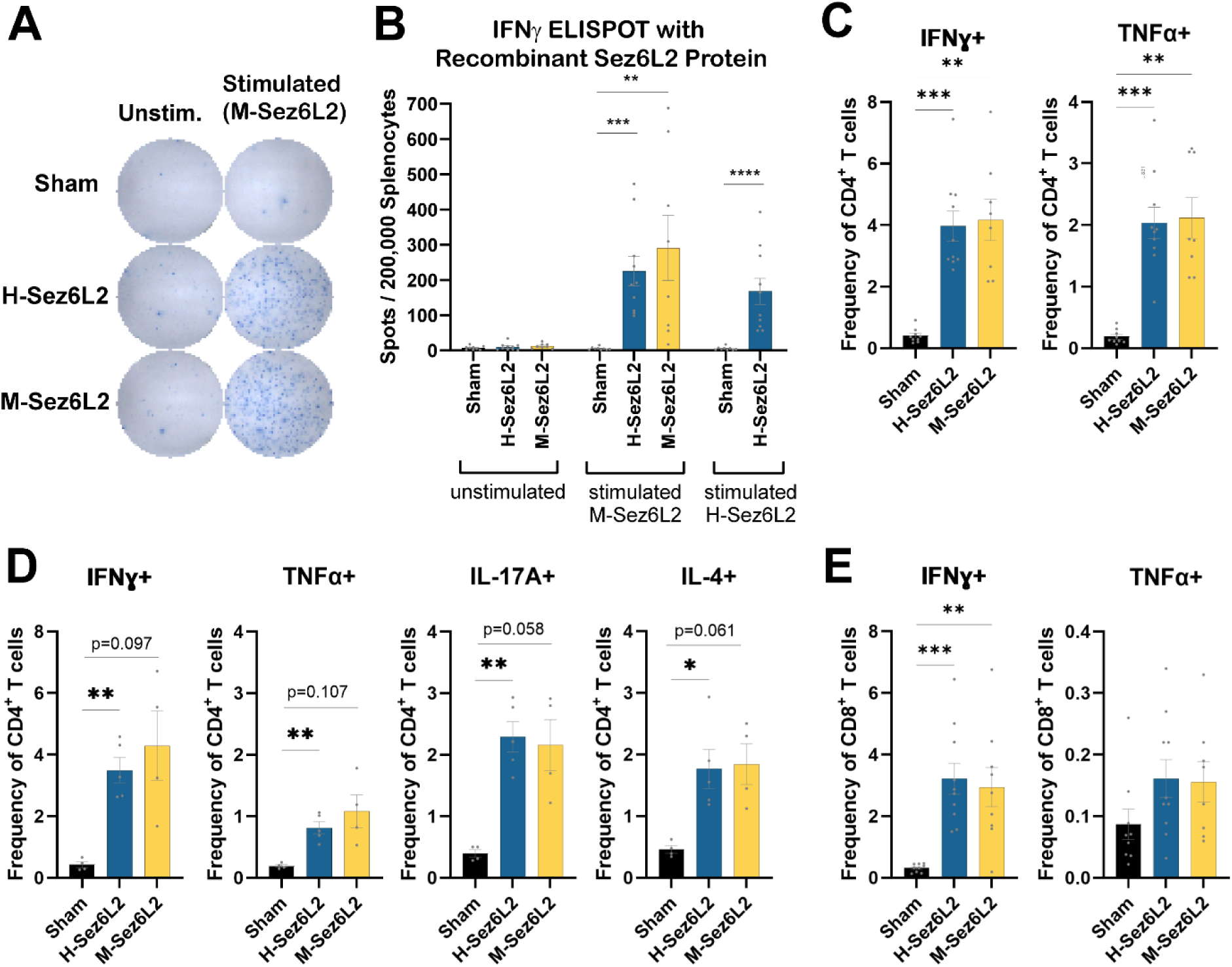
Sez6L2 immunized mice generate Sez6L2-specific T cells. A and B) Splenocyte/Lymph node mixed cell cultures from Sez6L2 or sham immunized mice were left unstimulated or were stimulated with H-Sez6L2 or M-Sez6L2 protein as indicated for 24 hours in an IFNγ ELISPOT assay. N=8-10. A) Representative images of ELISPOT wells. B) Quantification of the number of spots per 200,000 splenocytes/lymph node cells plated. C-E) Splenocyte/Lymph node mixed cell cultures from Sez6L2 or sham immunized mice were stimulated with M-Sez6L2 protein for 24 or 72 hours followed by cell surface and intracellular cytokine staining and analysis by flow cytometry. C) Graphs show the percent of CD4^+^ cells positive for IFNγ or TNFα after stimulation in culture for 24 hours. N=8-10. D) Graphs show the percent of CD4^+^ cells positive for IFNγ, TNFα, IL-4, or IL-17A after stimulation in culture for 72 hours. M=4-5. E) Graphs show the percent of CD8^+^ cells positive for IFNγ, TNFα after stimulation in culture for 24 hours. N= 8-10. Statistics for all graphs = Welch-ANOVAs with Dunnett’s T3 MCTs.

Having confirmed that Sez6L2-specific T cells were generated in our immunization model, we sought to identify the Sez6L2 immunodominant T cell epitopes. We designed a 15mer peptide library with a 10 amino acid overlap that contained peptides from both the human and mouse Sez6L2 proteins. Because we immunize mice with human Sez6L2 recombinant protein but are primarily interested in the autoimmune response occurring toward mouse Sez6L2, this combined human/mouse library allows us to assay the “autoimmune” peptide epitopes and the “foreign” peptide epitopes in the same assay. We immunized a new mixed-sex cohort of C57BL/6J mice with human Sez6L2 protein and compared them to a sham immunized group. This cohort also displayed motor impairments as they had more foot faults on the wire grid test and took more time to descend a rough metal pole (Figure 4 A, B). At 6 weeks post-immunization, we stimulated splenocytes with single peptides from our 15mer peptide library for immunodominant peptide responses using IFNɣ ELISPOT. We tested 188 total peptides in addition to the full recombinant mouse Sez6L2 protein (Figure 4 C, D). We found one immunodominant peptide that matched both mouse and human Sez6L2 (ATLGR<u>IVSPEPGGA</u>V) and one peptide that was unique to human (PS<u>WNGETPSCM</u>ASCG). We compared these findings to the MHC-II binding predictions made for H-Sez6L2 and the C57BL/6 H2-Ib1 allele by the NetMHCIIpan (ver 4.1) tool in the Immune Epitope Database (IEDB) Analysis Resource (40). Peptides with the IVSPEPGGA 9 amino acid core sequence were given the highest binding scores by NetMHCIIpan (Supplemental Figure 2). However, there were about 6 other core peptide sequences that received moderate prediction scores that did not show any hits in our ELISPOT screen. Similarly, our positive peptide that is unique to the human sequence containing the <u>WNGETPSCM</u> core sequence was given a low binding prediction score despite being biologically important. Our results show that immunization with the human Sez6L2 protein can achieve cross-reactivity towards the endogenous mouse Sez6L2. While the T cell activity directed at the “human only peptides” may be superfluous to a cell-mediated autoimmune response, these “human-directed” T cells may support autoimmunity in our model if they facilitate mouse Sez6L2-specific B-cell activation that drives production of autoantibodies.

**Figure 4:**
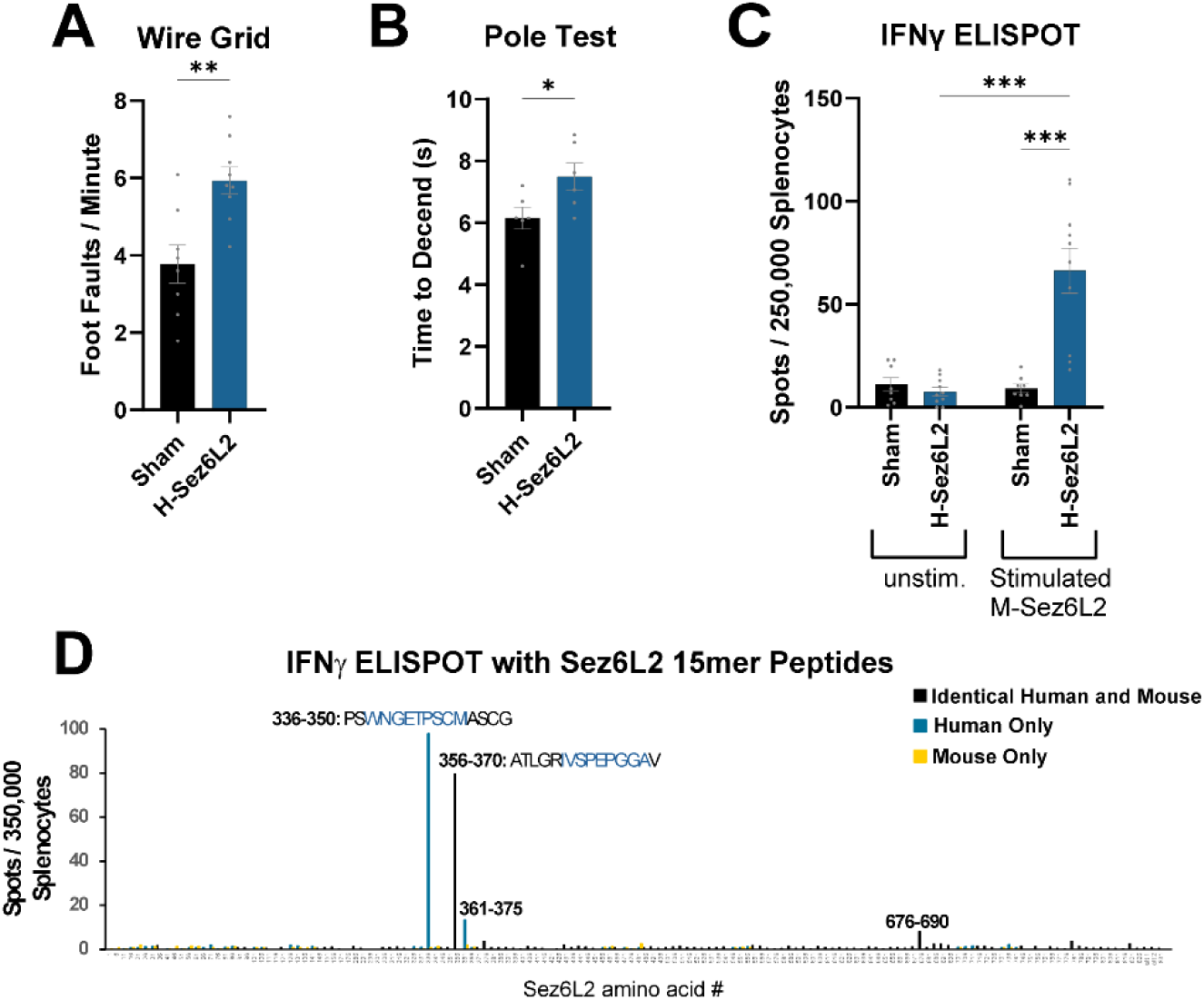
Immunodominant T cell epitopes from Sez6L2-immunized mice. A & B) Motor functions assessed at 5.5 weeks post-immunization on a mixed-sex cohort. Sham N=8; H-Sez6L2 N=9. A) Foot faults per minute of mobile activity on a wire grid. B) Time to descend a metal pole. Statistical tests: t test. C and D) Splenocytes from H-Sez6L2 or sham-immunized mice were stimulated with recombinant mouse Sez6L2 protein (C) or a 15mer/10aa overlapping peptide library containing peptides from the extracellular domain of mouse and human Sez6L2 (D) and assayed in an IFNγ ELISPOT assay. In D, the 15mer immunodominant peptide sequences are shown with the MHC-II core binding region predicted by IEDB Resource highlighted in blue. For the peptide ELISPOT, N=5 Sez6L2-immunized and N=3 for sham.

We next examined leukocyte infiltration in the brains of Sez6L2-immunized mice using flow cytometry. CD4+ T cell populations were approximately two-fold higher in the brains of H-Sez6L2- immunized mice compared to sham controls. Similarly, M-Sez6L2-immunized mice showed a trend toward increased CD4+ T cells in the brain, though this increase was not statistically significant. The frequencies of CD8+ T cells and CD19+ B cells in C57BL/6 brains remained unchanged (Figure 5 A).

**Figure 5:**
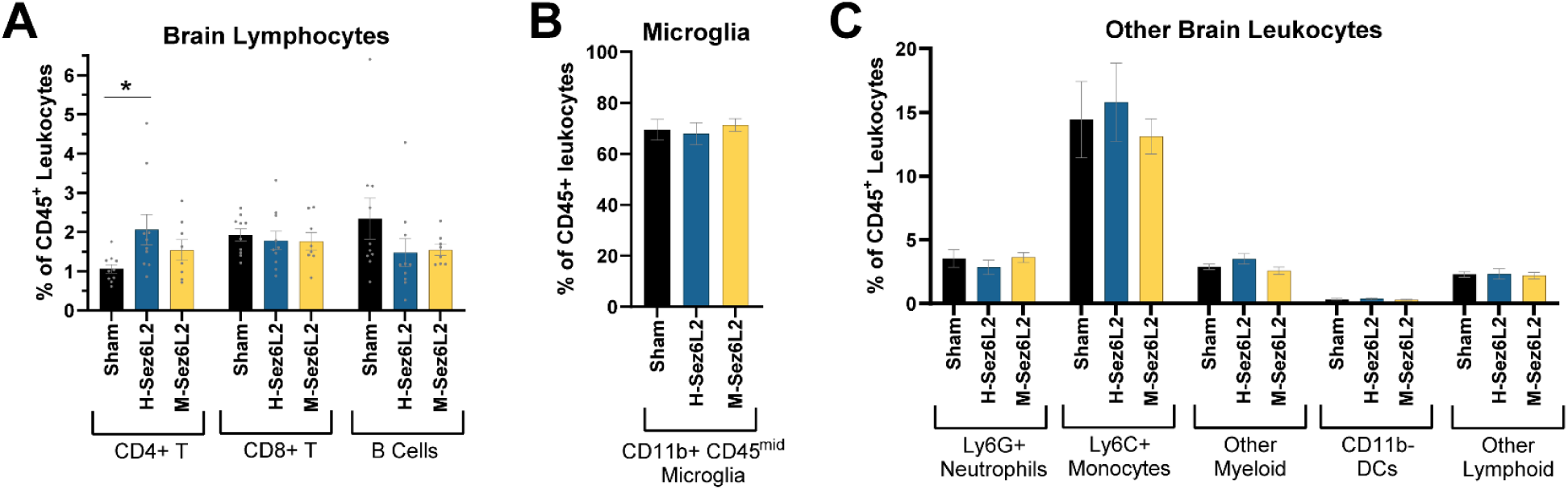
Mice Immunized with Sez6L2 have increased CD4^+^ T cell infiltration into the brain. A-C) Flow cytometry analysis of CD45+ immune cell populations in the brain. All cell populations are quantified as % of CD45^+^ cells. Statistics: Kruskal-Wallis Test with Dunn’s MCT for each cell type. Sham: n=10: 5 Male, 5 Female; H-Sez6L2: N=10: 5 Male, 5 Female; M-Sez6L2: N=8: 4 Male, 4 Female.

Additionally, quantification of other CD45+ cell populations revealed no changes in microglia, neutrophils, monocytes, or other minor leukocytes (Figure 5 B, C). The flow cytometry panel used in these experiments included markers for CD45, CD4, CD8, CD19, CD11b, LY6G, LY6C, and CD11C.

Finally, we evaluated the humoral immune response to Sez6L2 in immunized mice. ELISA analysis revealed significant levels of anti-mouse Sez6L2 antibodies in the serum of mice immunized with either H-Sez6L2 or M-Sez6L2 when tested at 6 weeks post-immunization (Figure 6A). Notably, H- Sez6L2-immunized mice exhibited higher levels of antibodies targeting mouse Sez6L2 at three weeks post-immunization compared to M-Sez6L2-immunized mice. This suggests that immunization with the human Sez6L2 protein induces an early, sustained, and robust humoral cross-reactivity against endogenous mouse Sez6L2. Using immunohistochemistry, we observed that the majority (5 of 7) of H- Sez6L2-immunized mice exhibited endogenous IgG localized to small subregions of the cerebellar parenchyma (Figure 6 B). Presumably, these endogenous IgGs are anti-Sez6L2 antibodies. The patchy distribution of the IgG suggests localized blood-brain barrier (BBB) disruption rather than widespread BBB or brain-CSF barrier (BCSFB) breakdown. In some mice, the endogenous IgGs were concentrated on Purkinje cell bodies, which are known to highly express Sez6L2. Additionally, reactive microglia (IBA1+ cells) were detected in and around cerebellar regions containing high IgGs, indicating that infiltrating autoantibodies may induce neuroinflammation (Figure 6 B).

**Figure 6:**
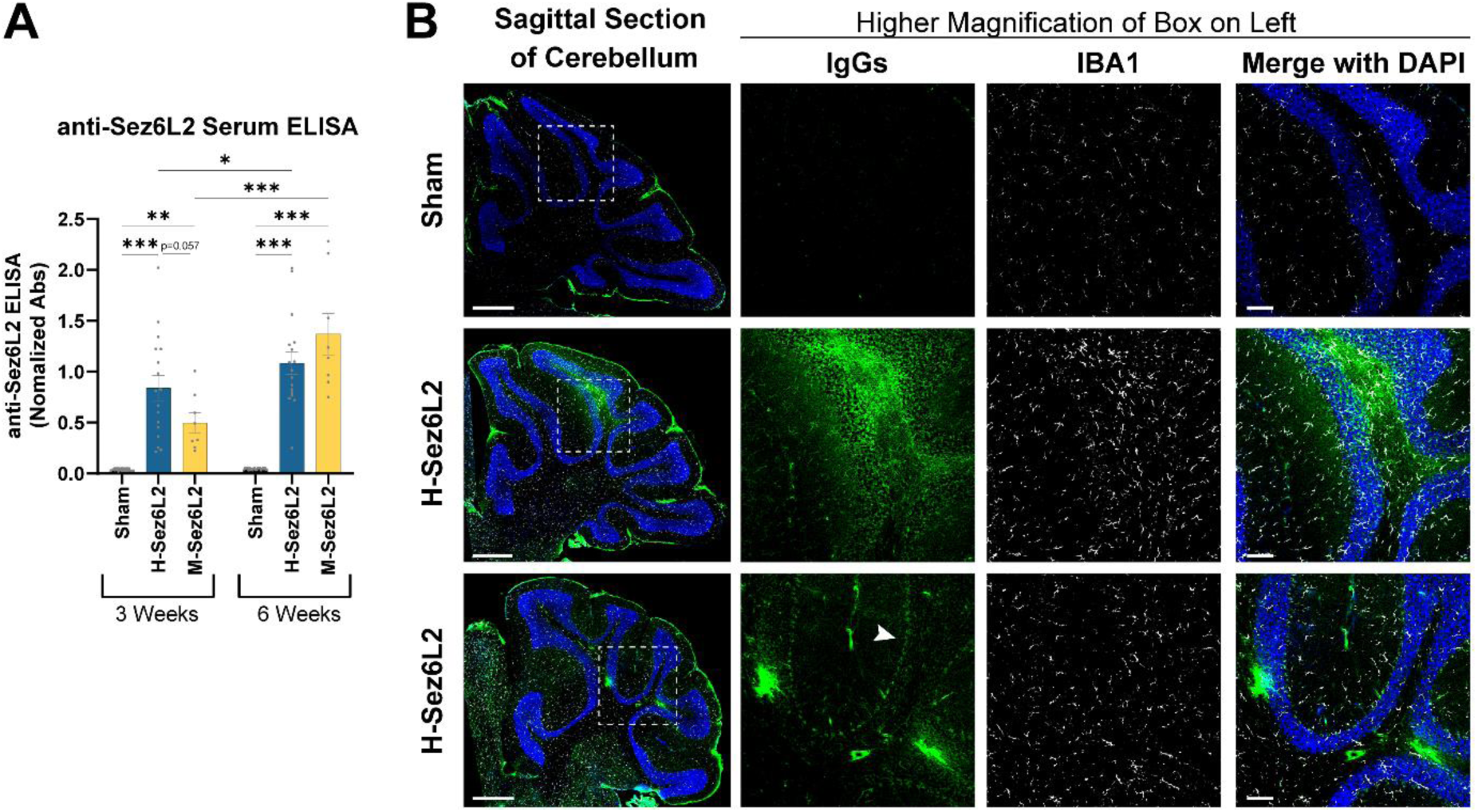
Sez6L2 immunized mice generate significant anti-Sez6L2 antibodies that can infiltrate into the cerebellar parenchyma. A) An anti-Sez6L2 ELISA was used to detect Sez6L2 antibodies in serum collected from immunized mice at 3- and 6-weeks post-immunization. Sham: n=22: 11 Male, 11 Female; H-Sez6L2: N=17: 9 Male, 8 Female; M-Sez6L2: N=8: 4 Male, 4 Female. B) IHC for endogenous IgGs (green), IBA1^+^ microglia/myeloid cells (white) and DAPI (blue) using cerebellar sections from immunized mice collected at 6 weeks post-immunization. Endogenous IgGs are limited to the meninges in sham mice. Sez6L2-immunized mice have small regions of the cerebellar parenchyma where high levels of endogenous IgGs have infiltrated. They also exhibited areas with more difuse IgGs bound to Purkinje cell bodies. Two representative H-Sez6L2 samples are shown to illustrate the various patterns of IgG infiltration. Increased IBA1 fluorescence in the areas of IgG infiltration suggest Sez6L2 autoantibodies induced reactive microglia.

## DISCUSSION

Autoimmune diseases of the central nervous system are increasingly recognized for their complex pathophysiology and diverse clinical manifestations. This study demonstrates that an autoimmune response to Sez6L2 in mice can result in mobility impairments and pathology resembling that observed in human patients with cerebellar ataxia associated with Sez6L2 autoantibodies (3–14). Our findings revealed that Sez6L2-immunized mice have motor impairments quantified through open field, wire grid walk, and pole test assays. Furthermore, these mice developed significant levels of anti- Sez6L2 antibodies and Sez6L2-specific T cells. Sez6L2-immunized mice also showed the presence of IgGs and reactive microglia in the cerebellar parenchyma, along with increased CD4+ T cell infiltration into the brain, supporting the hypothesis that Sez6L2 autoimmunity contributes to neurological injury.

The detection of anti-Sez6L2 antibodies in the serum and their localization within the cerebellar parenchyma suggest that antibody-mediated mechanisms play a significant role in the pathology of Sez6L2 autoimmunity. As Sez6L2 is expressed on the cell surface, these antibodies can directly bind neurons potentially leading to cell damage and dysfunction through mechanisms observed in other autoantibody-mediated diseases, including complement activation, antibody-dependent cytotoxicity (ADCC), opsonization and cellular phagocytosis, or direct impairment of Sez6L2 function (42). The presence of reactive microglia in regions with IgG deposition further supports the role of antibody-mediated neuroinflammation in localized cerebellar damage.

In addition to antibody-mediated mechanisms, our findings underscore the importance of CD4+ T cell-mediated pathology in Sez6L2 autoimmunity. The increased infiltration of CD4+ T cells into the brain suggests that these cells contribute to disease progression and the creation of an inflammatory environment. The generation of multiple CD4+ T cell subtypes (Th1, Th2, and Th17), along with putative CD8+ T cells capable of targeting endogenous Sez6L2, highlights a complex and multifaceted immune response. In the experimental autoimmune encephalomyelitis (EAE) mouse model of multiple sclerosis, both Th17 and Th1 T cells are known to breach the BBB and BCSFB to enable immune cell entry into and injury of the central nervous system (reviewed in (43)). Similarly, Sez6L2-specific T cells in our model may play a critical role in breaching the BBB, thereby allowing immune cell and antibody access to the cerebellum and other brain regions. Understanding the relative contributions of antibody-mediated versus T cell-mediated pathology is essential for developing effective treatments for patients and remains an ongoing focus of our work on Sez6L2 autoimmunity.

Our results show that immunizing mice with either human or mouse recombinant Sez6L2 induced production of Sez6L2 antibodies and Sez6L2-specific T cells capable of targeting mouse Sez6L2. However, immunization with human Sez6L2 alone led to significant motor deficits and produced a valuable mouse model of Sez6L2 autoimmunity. It is likely that the minor differences in protein sequence between human and mouse Sez6L2 helped bypass tolerance mechanisms and allowed sufficient species cross-reactivity to facilitate the autoimmune response. This is supported by our identification of two immunodominant T cell epitopes for the human Sez6L2 protein, including one shared between human and mouse Sez6L2 and one unique to human Sez6L2. Additionally, immunization with human Sez6L2 generated higher levels of Sez6L2 antibodies at the early 3-week timepoint compared to immunization with mouse Sez6L2. This early and sustained production of Sez6L2 antibodies in mice immunized with human Sez6L2 may explain why they developed significant motor impairments, whereas the mice immunized with mouse Sez6L2 did not.

Our study introduces a novel mouse model of Sez6L2 autoimmunity that replicates key features of cerebellar ataxia observed in human patients with Sez6L2 autoantibodies. This model offers a valuable platform for studying immune pathways involved in brain injury and for testing therapeutic strategies to modulate the immune response. By elucidating the mechanisms driving this disease, we aim to accelerate the development of effective treatments and improve outcomes for patients affected by this condition.

## Supplementary Material

Supplementary material is available online at journal’s site.

## List of Abbreviations

CSF: cerebral spinal fluid
BBB: blood brain barrier
BCSFB: blood CSF barrier.

## DECLARATIONS

### Ethics approval and consent to participate

Animal care and use were carried out in compliance with the US National Research Council’s Guide for the Care and Use of Laboratory Animals and the US Public Health Service’s Policy on Humane Care and Use of Laboratory Animals. Protocols were approved by the University Committee on Animal Resources at the University of Rochester.

### Consent for publication

Not Applicable

### Availability of data and materials

The datasets used and/or analyzed during the current study are available from the corresponding author on reasonable request.

### Authors’ contributions

JH designed the study. All authors help care for the Sez6L2 immunized mice and performed end-point tissue collection and processing. CRS, JR, and JH immunized the mice. CRS, JR, AR, and JH completed the behavior studies. CRS performed the Sez6L2-specific T cell flow cytometry assays. JG with the help of AH, AR, CRS, and JH performed the Elispot assays. JR performed the flow cytometry experiments on brain samples. CRS, JR, and AR performed the antibody ELISAs. CRS, JR, AR, HL, and JH processed, imaged, and analyzed the IHC experiments. All authors read and approved the final manuscript.

## Acknowledgements

The Genotype-Tissue Expression (GTEx) Project was supported by the Common Fund of the Office of the Director of the National Institutes of Health, and by NCI, NHGRI, NHLBI, NIDA, NIMH, and NINDS. The data used for the analyses described in this manuscript (Figure 1C) were obtained from: the GTEx Transcript Browser on the GTEx Portal on 06/21/24. We also thank Dr. Harris Gelbard, University of Rochester Medical Center, for his support and helpful feedback on the project.

