## Supplemental for "Sez6L2 autoimmunity induces cerebellar ataxia in mice"

Supplemental Figures and Legends:

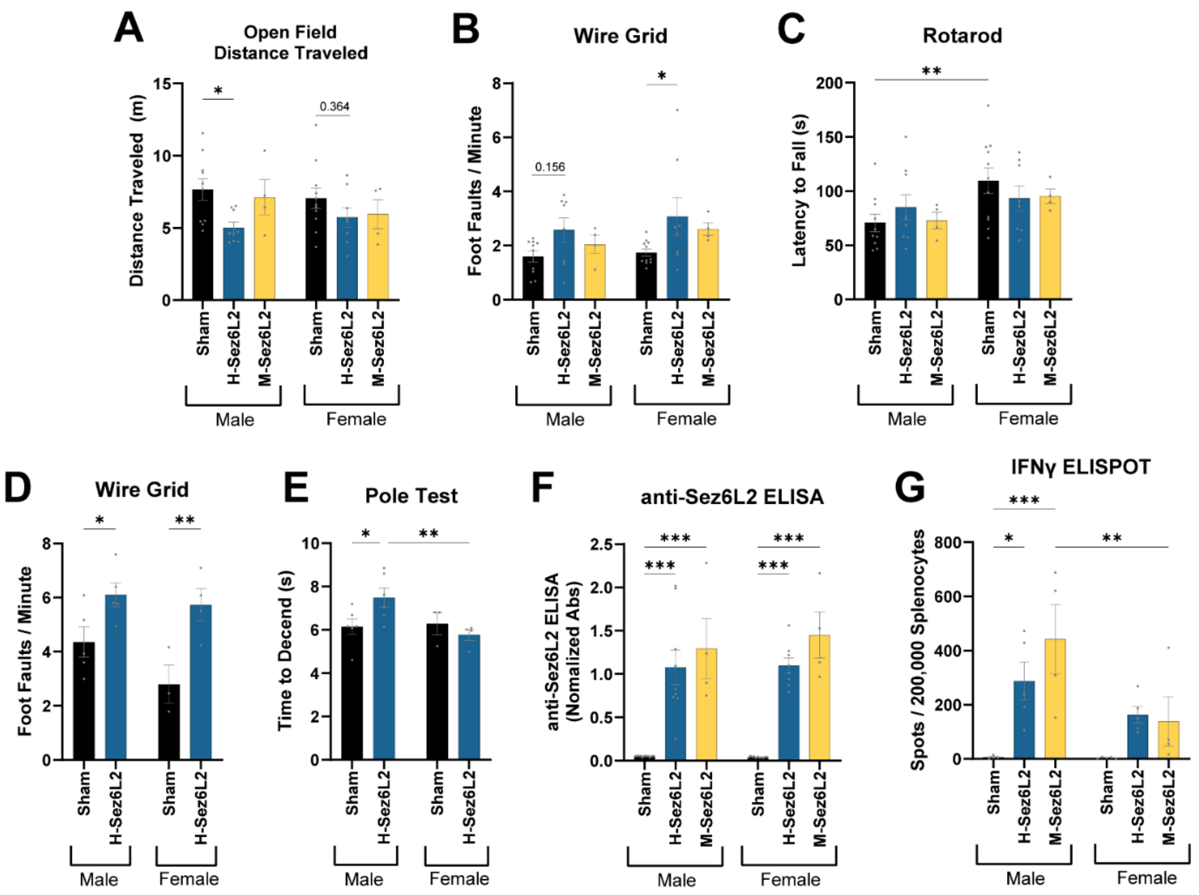


Supplemental Figure 1: Behavior, Sez6L2 antibody ELISA, and ELISPOT data separated by Sex. A) Cohort 1: Distance traveled in a 10-minute open field assay performed at 5.5-weeks post-immunization. B) Cohort 1: Foot faults per minute on wire grid performed at 5.5 weeks post-immunization. C) Cohort 1: Latency to fall off accelerating rotarod performed at 5.5 weeks post-immunization. D) Cohort 2: Foot faults per minute on wire grid performed at 5.5 weeks post-immunization. D) Cohort 2: Time to descend a pole at 5.5 weeks post-immunization. F) Cohort 1: Sez6L2 antibody ELISA was used to detect Sez6L2 antibodies in serum collected from immunized mice at 6-weeks post-immunization. G) Cohort 1: Splenocyte/Lymph node mixed cell cultures from Sez6L2 or sham immunized mice were left unstimulated or were stimulated with M-Sez6L2 protein for 24 hours in a IFNγ ELISPOT assay. Statistics for all graphs: 2-way ANOVA with Tukey's MCT.

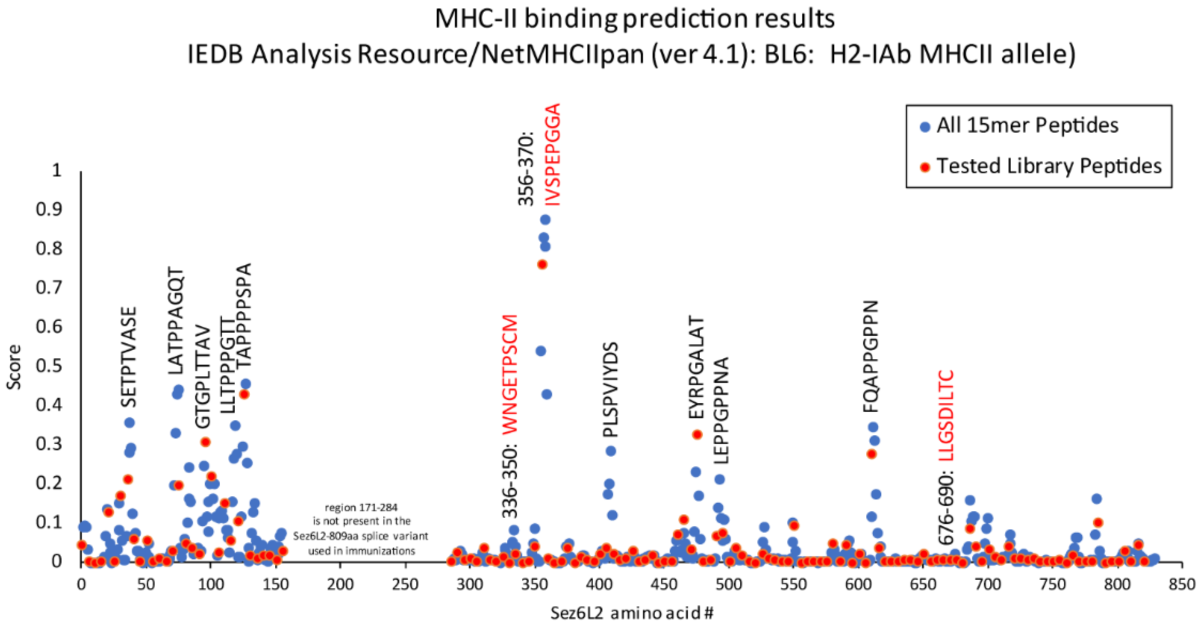


Supplemental Figure 2: Human Sez6L2 MHC-II binding prediction scores for the murine C57BL/6 H2-Ib1 allele by the NetMHCIIpan (ver 4.1) tool available from the Immune Epitope Database (IEDB) Analysis Resource. Blue dots represent scores for all 15mer peptides in the human Sez6L2 extracellular domain immunizing protein. Red dots represent the tested peptides used in our 10aa overlap library. Peptide sequence labels with red letters had positive ELISPOT readings in our assay; peptide sequence labels with black letters were negative in our ELISPOT assay but were given moderate scores by the NetMHCIIpan tool. Note: The Sez6L2 peptides are labeled according to the amino acid numbering of the canonical splice variant of Sez6L2 that is 910 amino acids in length even though we immunized with a shorter splice variant that was 809 amino acids in length.
